# Succinate metabolism uncouples retinal pigment epithelium mitochondria

**DOI:** 10.1101/2021.02.10.430650

**Authors:** Daniel T. Hass, Celia M. Bisbach, Brian M. Robbings, Varun S Kamat, Martin Sadilek, Ian R. Sweet, James B. Hurley

## Abstract

Succinate is central in energy metabolism. It is thought to be made and consumed within mitochondria, and not to cross the plasma membrane. Contrary to the canonical role of succinate, it is produced in the retina and released into the extracellular milleau. It is unknown how this affects adjacent retinal pigment epithelium, within eyecup tissue. By measuring consumption of ^13^C_4_-succinate and O_2_ we probe metabolism of the eyecup, we find robust succinate consumption by eyecup but not retina tissue. Succinate-dependant stimulation of eyecup respiration is oligomycin resistant; indicating that in eyecup tissue succinate uncouples mitochondrial ATP synthesis from respiration.

## Introduction

Succinate is a critical intermediate in the tricarboxylic acid cycle and electron transport chain. When oxidized to fumarate it fuels reduction of O_2_ to H_2_O. The canonical view of succinate only as an e- donor is incomplete, since some tissues are net producers of succinate. Cells in the retina ^1^, pancreas ^2^, exercising muscle ^3,4^, and in hypoxic tissues ^5,6^ can each export succinate.

Succinate formed in these tissues may be released into the extracellular space or into circulation. Some tissues such as brown adipose tissue are capable of using circulating succinate to fuel thermogenesis ^7,8^, yet overall succinate turnover in circulation is thought to be low ^9^. This may be because succinate is not a metabolic fuel in most tissues, and indeed some studies have argued that succinate does not cross the plasma membrane ^10–12^. Alternately in some tissue niches succinate may be both generated and consumed before reaching circulation, which would require a robust ability for succinate uptake.

Our past work to understand succinate metabolism in the eye revealed that retinas release succinate and eyecup (composed of choroid, sclera, and retinal pigment epithelium) OCR is responsive to succinate ^1^. We further investigated succinate metabolism in tissues of the eye and surprisingly found not only a robust capacity for intact cells to take up succinate but also that oxidation of succinate by eyecup tissue stimulates mitochondrial uncoupling.

## Results

Within cells, succinate oxidation to fumarate stimulates O_2_ consumption (OCR) by complex IV. If succinate is tissue-impermeant, tissues should not deplete it from media and it should not stimulate OCR. In line with this hypothesis, succinate minimally affects retina tissue OCR over a wide range of concentrations (**Figure 1B**). The same succinate concentrations strongly stimulate OCR in eyecup tissue (containing retinal pigment epithelium, choroid, and sclera) (**Figure 1A**). Similarly, depletion of ^13^C_4_-succinate from culture medium by eyecups was strongly concentration dependent (**Figure 1C**). As in our past experiments, these data suggest that succinate oxidation is limited in the retina and robust in the eyecup ^1^.

**Figure 1.**
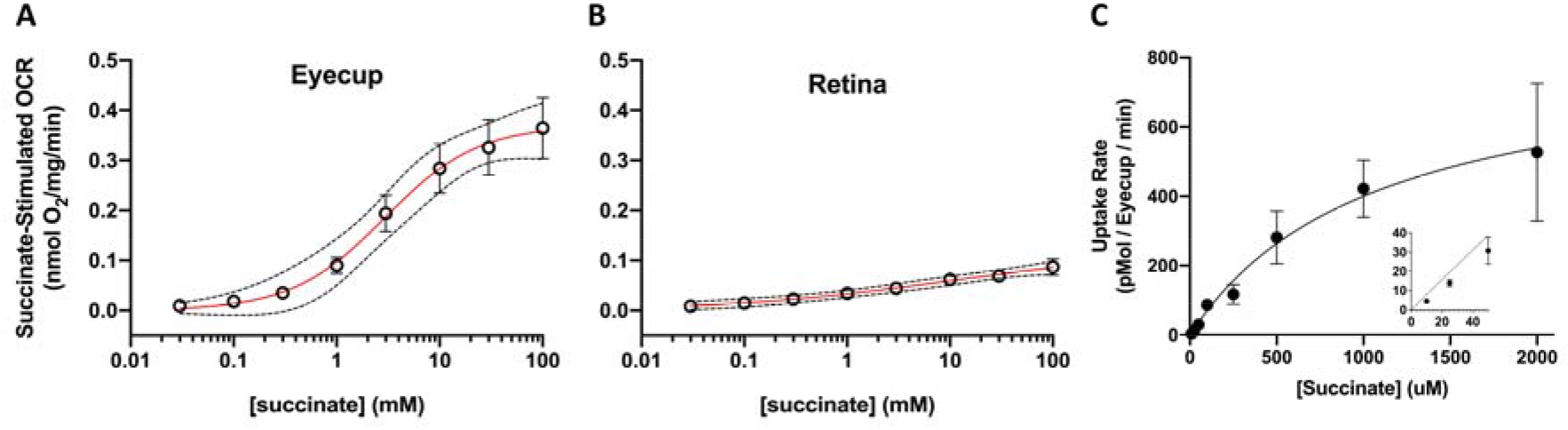
Extracellular Succinate Selectively Increases Oxygen Consumption in Ocular Tissues. We measured O_2_ consumption rate (OCR) in freshly dissected (**A**) eyecup (n=3) and (**B**) retina tissue from C57BL6/J mice, respiring in KRB buffer with 5 mM glucose. We supplied this media with increasing concentrations of disodium succinate (30 μM, 100 μM, 300 μM, 1 mM, 3 mM, 10 mM, 30 mM, or 100 mM) and determined the mean ± SEM steady state OCR above the 5 mM glucose “baseline” respiration. We fit the data with an allosteric sigmoidal curve (red lines). Best-fit parameters are available in **Table 1**. Dotted lines surrounding the curves represent 95% confidence intervals from the curve fit. (**C**) Mean ± SEM succinate uptake rate from eyecup medium were also determined.

Succinate-stimulated OCR could reflect a biological capacity for succinate uptake and oxidation, or alternately the possibility that eyecup isolation damages plasma membranes of cells within the tissue. We used two distinct methods (assessment of ^14^C-sucrose uptake and measurement of LDH release) to determine if our tissue preparations contain intact cells. We first measured permeability by assessing uptake of ^3^H_2_O and ^14^C-sucrose uptake in retina and eyecup tissue. There is no known sucrose transporter in mice, so intact tissue should exclude ^14^C-sucrose but not ^3^H_2_O. We compared ^3^H2O or ^14^C-sucrose uptake after an hour in culture, and quantified uptake as a % of total radioactive dose. Eyecup and retina tissue retained 0.34% and 0.17% of the ^3^H2O dose, and 0.09% and 0.05% of the ^14^C-sucrose dose. We determined tissue permeability by calculating the ratio of radioisotope uptake, though this may overestimate ^14^C-sucrose if it is present in eyecup vasculature and other extracellular compartments. This measure suggests that retinas are 25.4±0.9% sucrose-permeable and eyecups are 28.6±2.3% sucrose-permeable (mean ± SEM, **Figure 2A**).

**Figure 2.**
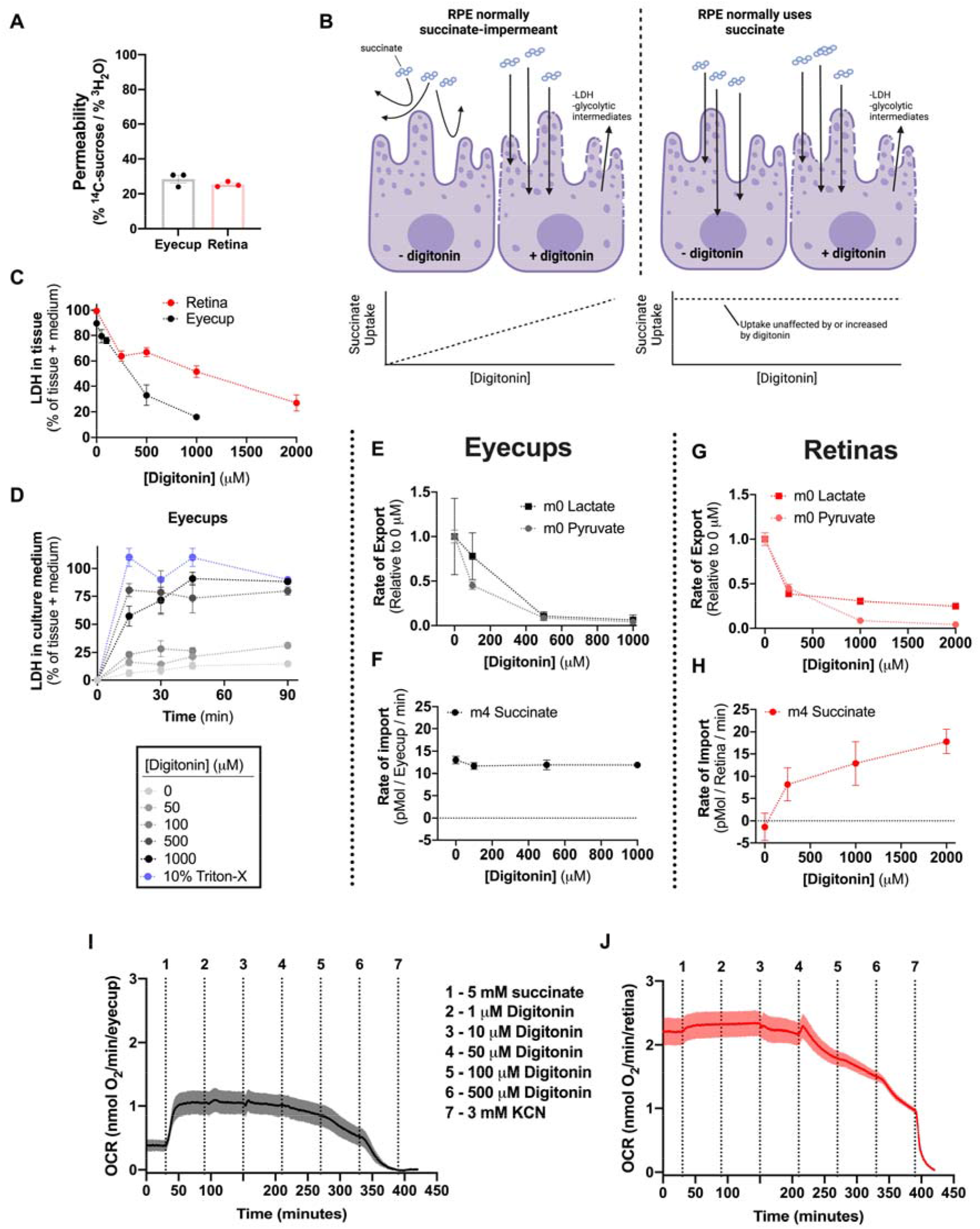
RPE Permeability does not alter Succinate Oxidation. (**A**) Permeability of freshly dissected retina and RPE tissue assessed by the relative uptake of ^3^H_2_O and ^14^C-sucrose over an hour *ex vivo*. (**B**) Model of the hypothesis to test whether succinate oxidation by eyecup tissue is the result of cell permeability. (**C**) Retina and eyecup [LDH] increases as a function of [digitonin]. Percentage of total was determined by assaying LDH in both tissue and culture medium when [LDH] was at a steady state in medium. (**D**) Determination of the *ex vivo* culture time needed to reach steady-state [LDH]. (**E**) Relative lactate and pyruvate release rate by eyecups decreases with increasing [digitonin]. (**F**) Eyecup ^13^C_4_-succinate uptake rate however is unaltered by [digitonin]. (**G**) As with eyecup tissue, lactate and pyruvate release decreases in retinas with increasing [digitonin]. (**H**) Unlike with eyecups, retina ^13^C_4_-succinate uptake increases with [digitonin], suggesting that the plasma membrane is a barrier for retina but not eyecup succinate uptake. (**I**) This was corroborated by measurements of eyecup and (**J**) retina O_2_ consumption rate before and after addition of 5 mM succinate and increasing concentrations of digitonin. Digitonin did not enhance eyecup OCR in the presence of succinate, and OCR only fell with higher concentrations of digitonin.

If membranes are broken by dissection, cell components such as lactate dehydrogenase (LDH) and metabolites will diffuse into culture medium over time, which should disable normal metabolism (**Figure 2B**). We tested this hypothesis and found minimal LDH release from retina (∼0% of tissue LDH) or eyecup tissue (<10% of tissue LDH) over an hour of culture. To show that increased permeability leads to LDH and metabolite leakage we used increasing concentrations of the non-ionic detergent digitonin as a positive control (**Figure 2C**) and confirmed that LDH release had reached a steady state by measuring enzyme concentrations in medium aliquots over time (**Figure 2D**). Taken together the radioactivity and LDH release experiments show that cells in freshly dissected retinas and eyecup tissue are mostly intact.

Eyecups do however contain permeabilized cells. Several groups have demonstrated that extracellular succinate is not oxidized well by intact cells and we next tested the population of permeabilized cells supports the the succinate oxidation we have observed. If this occurs, increasing cell permeability should allow a greater portion of cells to access succinate, increasing OCR and ^13^C_4_-succinate uptake. We used gas chromatography-mass spectrometry (GC-MS) to assess metabolite uptake and release rates in control and permeabilized tissue. We sampled media from tissues incubated in 5 mM glucose, 50 μM ^13^C_4_-succinate, and increasing concentrations of digitonin. Digitonin decreased release of the glycolytic end products lactate and pyruvate (**Figure 2E,G**), showing that both in retina and eyecup tissue, glucose metabolism is short-circuited by nonselective membrane permeability, likely resulting in the diffusion of glycolytic enzymes or early glycolytic intermediates or out of the tissue.

In eyecup tissue, digitonin did not affect the rate of ^13^C_4_-succinate uptake (**Figure 2F**). In retinas, digitonin increased ^13^C_4_-succinate uptake, suggesting that plasma membrane permeability normally blocks retina but not eyecup succinate use (**Figure 2H**). Succinate uptake should stimulate mitochondrial activity, and we tested this hypothesis by measuring OCR in the presence of succinate, before and after adding in 1, 10, 30, 100, and 500 μM digitonin (**Figure 2I**). None of these concentrations increased retina or eyecup OCR. Rather, increasing digitonin concentration suppressed respiration in both tissues. The decrease in respiration results either from a direct interaction between digitonin and the electron transport chain or because in a sufficiently permeabilized tissue preparation the flow of medium carries dislodged metabolites or mitochondria away in the medium outflow. These experiments show that eyecup plasma membrane permeability does not scale with succinate uptake or oxidation. We also show that retina succinate consumption is possible only in the presence of a permeabilizing agent, but that succinate consumption in the presence of digitonin likely cannot match the energetic contribution that glucose oxidation makes to the retina.

Succinate stimulates eyecup respiration more drastically than many other substrates ^1^.,Succinate is a direct substrate for the ETC other metabolites we have tested in the past (lactate, glutamine, proline) require several sequential reactions to produce NADH used by the ETC. To determine if succinate-stimulated oxygen consumption is a unique phenomenon or due only to the direct nature of succinate as an ETC fuel, we tested the hypothesis that other more direct mitochondrial substrates (equimolar concentrations of pyruvate and malate) could influence respiration similarly to succinate. Pyruvate and malate together should sustain OCR through NADH. We compared OCR as a function of substrate concentration and found pyruvate and malate do not stimulate OCR to the same levels as succinate (**Figure 3A**).

**Figure 3.**
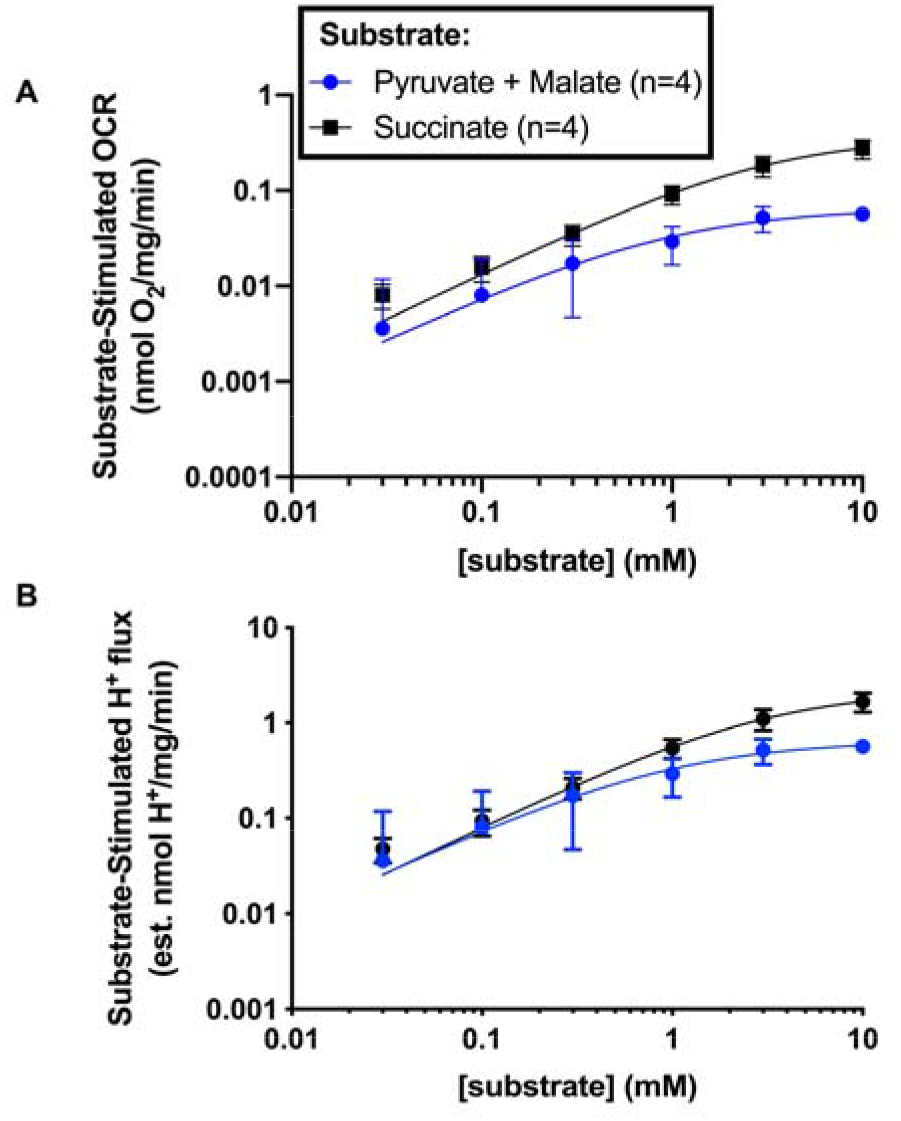
Succinate is a preferred substrate for eyecup mitochondria **(A)** Eyecup OCR respiring on 5 mM glucose supplemented with increasing concentrations (30 μM, 100 μM, 300 μM, 1 mM, 3 mM, 10 mM) of succinate or malate and pyruvate. (**B**) We estimated mitochondrial H^+^ flux from the matrix to the intermembrane space by multiplying OCR by the H^+^/2e^-^ ratio for the pyruvate/malate-linked substrate NADH (10) or succinate (6) ^18^.

This comparison however may not fully represent the ability of each substrate to generate ATP for two reasons. The first is that each molecule of succinate reduces a single FADH_2_ before exiting mitochondrial metabolism, but each pyruvate/malate pair forms three NADH molecules before exiting mitochondrial metabolism. We respectively show these abbreviated TCA cycle activities diagrammed (**Figure 3B**) and through an analysis of U-^13^C-succinate and U-^13^C-pyruvate flux in eyecup tissue (**Figure 3C-D**), We can adjust for this difference in electron input by tripling the [substrate] when eyecups respire on pyruvate/malate. In addition, NADH- and succinate-linked respiration each translocate a different number of H^+^ for each O_2_ molecule consumed. To compare the driving force for ATP production with succinate or pyruvate/malate, we converted substrate-dependent OCR into estimates of H^+^ flux, assuming that each O_2_ molecule consumed using NADH linked substrates catalyzes translocation of 10 H^+^, while succinate-dependent OCR translocates 6 H^+^ (**Figure 3E**). With this correction the effect of substrate concentration on H^+^ translocation overlapped at ≤300 μM but differed at ≥1 mM. Above 1 mM, succinate cause more H^+^ flux than equimolar pyruvate/malate.

The mismatch in H^+^ flux suggests that exogenous succinate produces a H^+^ leak across the inner mitochondrial membrane, and the rate of substrate oxidation increases to compensate for the lost driving force for ATP production caused by H^+^ leak (**Figure 4A**). We determined whether succinate produces H^+^ leak by exposing eyecup tissue to either 5 mM succinate or 5 mM pyruvate/malate in the presence of the ATP synthase inhibitor oligomycin. O_2_ consumption in the presence of oligomycin represents substrate oxidation that is not inhibited by the mitochondrial proton-motive force (Δρ). With glucose alone oligomycin completely suppresses OCR, indicating that when eyecup mitochondria respire using glucose, ATP synthesis and e^-^ transport are coupled. Adding 5 mM of either substrate to this cocktail increased eyecup respiration though the effect of succinate is several fold greater than that of pyruvate/malate (**Figure 4B**). The effect of [succinate] on eyecup OCR is surprisingly intact in the presence of oligomycin over a broad range of concentrations (**Figure 4C**). Rotenone blocks a portion of succinate-driven OCR, suggesting that succinate is able to re-activate NADH oxidation, which was previously suppressed by oligomycin (**Figure 4D**). These data suggest that OCR with succinate as a substrate is high because succinate introduces an oligomycin-insensitive H^+^ ‘leak’ to otherwise well-coupled mitochondria. This leak enables rapid succinate oxidation and allows for additional NADH-dependent respiration when it has previously been inhibited by a steep H^+^ gradient.

**Figure 4.**
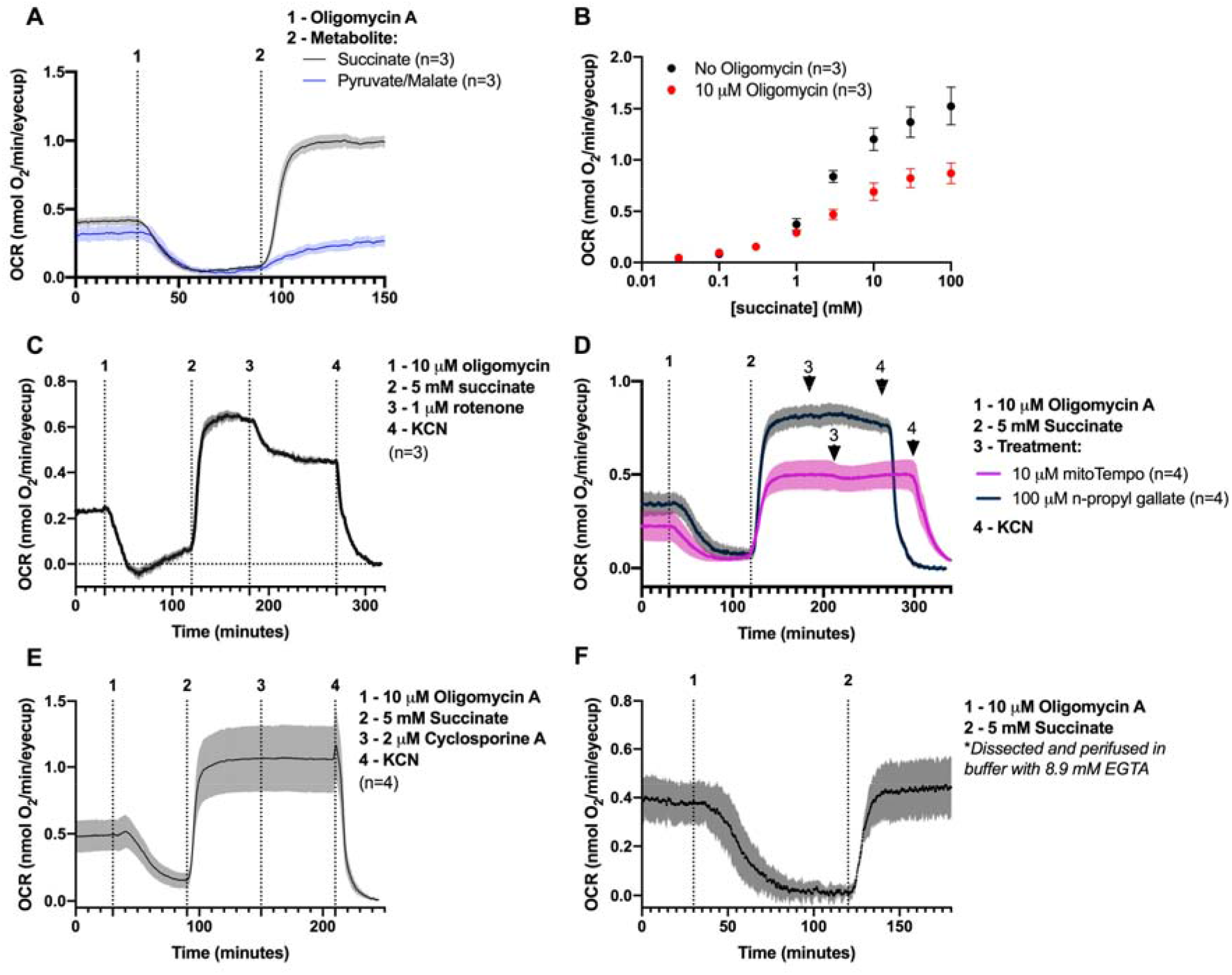
Extracellular Succinate Uncouples *ex vivo* Eyecup Mitochondria. (**A**) We determined ATP synthase-independent substrate oxidation by incubating eyecup tissue in 5 mM glucose, then in 5 mM glucose with 10 μM of the ATP-synthase inhibitor oligomycin, and finally in a mix of 5 mM glucose, 10 μM oligomycin, and 5 mM of succinate (black) or pyruvate/malate (blue). (**B**) We determined OCR as a function of [succinate] in the presence or absence of 10 μM oligomycin A. (**C**) To determine whether NADH oxidation was the source of uncoupled respiration, we first stimulated succinate-dependent uncoupling as in A, then added 1 μM of the complex I inhibitor rotenone. Which partially inhibited respiration, suggesting that succinate re-actives NADH oxidation after oligomycin shuts it down. To determine the source of oligomycin-resistant respiration, we performed the treatments described in (A), but following the addition of 5 mM succinate we attempted to shut down the succinate-dependent increase in OCR by adding (**D**) the antioxidants mitoTempo (purple line) or n-propyl gallate (black line), (**E**) the mitochondrial permeability transition pore complex inhibitor cyclosporine A, or (**H**) EGTA to deplete perifusion medium of free Ca^2+^ during the experiment. Lines represent mean O_2_ consumption rate following subtraction of the 5 mM glucose “baseline” (**B**) or the 3 mM KCN “floor” (**A**,**C-F**).

Multiple mechanisms could cause a H^+^ leak, including reactive oxygen species (ROS)-driven activation of an uncoupling protein ^7,13^ or activation of the mitochondrial permeability transition pore complex (mPTP) ^14^. If succinate increases [ROS] and ROS drive mitochondrial uncoupling, antioxidants should be sufficient to decrease or prevent H^+^ leak. Neither 50 μM mitoTempo or 100 μM n-propyl gallate clearly affected succinate-driven OCR (**Figure 4E)**. If the mPTP was responsible, inhibiting the mPTP with cyclosporine A (**Figure 4F**) or sequestering free Ca^2+^ with 8.6 mM EGTA (**Figure 4G**) should prevent oligomycin-insensitive OCR. However none of these conditions suppressed succinate-dependent leak respiration.

## Discussion

Our study has revealed two main novel concepts that may be important in the biology of the eye. The first is that intact eyecup tissue takes up and oxidizes succinate. We show this by measuring oxygen consumption and ^13^C_4_-succinate uptake. Eyecup tissue is minimally permeabilized after dissection, and neither OCR nor tracer uptake is enhanced with higher cell permeability (**Figure 2**). This finding is consistent with a report that succinate stimulates consumption of O_2_ by explants of dog heart and rabbit kidney ^15^. However, other groups have reported that succinate does not stimulate metabolism in most cultured cells ^7,11,12^. We believe these data together reflects an unappreciated degree of cell-specificity to succinate uptake.

Our second main finding is that eyecup mitochondria gain a H^+^ leak (become uncoupled) when succinate is a metabolic substrate (**Figure 4**). We show this through an increase in oligomycin-resistant respiration. Uncoupling with succinate appears to re-enable NADH oxidation and drastically affects OCR in the eyecup.

Why does succinate uncouple eyecup mitochondria? The answer is unclear, but succinate oxidation is capable of stimulating a high Δρ and in isolated mitochondria there is an exponential relationship between Δρ and H^+^ leak. The current best explanation for this relationship is that at high Δρ-dependent there is a dielectric breakdown of the mitochondrial membrane ^16,17^. Follow-up investigations should be focused on determining if this is truly the mechanism of H^+^ leak in eyecup tissue.

To our knowledge this is the first report of succinate-dependent uncoupling in intact tissue. In a previous study we established the possibility that eyecup tissue may oxidize succinate from the retina ^1^, and our new data add more context to the model of succinate and malate exchange we proposed to normally occur between retina and RPE. If the retina is constantly releasing succinate, mitochondria in the eyecup may be uncoupled regularly. If that occurs, it may explain the surprisingly low O2 levels in regions of the retina near the RPE. Low retinal [O2] is thought to be a consequence of high OCR in the retina, but our data show comparable or even greater OCR in the succinate-stimulated eyecup. This suggests the possibility that in vivo, eyecup tissue may contend with the retina as a main O_2_ sink.

## Materials and Methods

### Ethical Approval

This study was carried out in accordance with the National Research Council’s Guide for the Care and Use of Laboratory Animals (*8th ed*). All protocols were approved by the Institutional Animal Care and Use Committees at the University of Washington.

### Animals

All experiments used 2-5 month-old male and female wild-type C57BL6/J mice. These mice were housed at an ambient temperature of 25°C, with a 12-hour light cycle and *ad libitum* access to water and normal rodent chow.

### Ex vivo metabolic flux

In all *ex vivo* labeling experiments, we quickly euthanized mice by awake cervical dislocation, dissected the indicated tissues in Hank’s Buffered Salt Solution (HBSS; GIBCO, Cat#: 14025-076), and incubated them in pH 7.4 Krebs-Ringer bicarbonate (KRB) buffer (98.5 mM NaCl, 4.9 mM KCl, 1.2 mM KH_2_PO_4_ 1.2 mM MgSO_4_-7H_2_O, 20 mM HEPES, 2.6 mM CaCl_2_-2H_2_O, 25.9 mM NaHCO_3_) supplemented with 5 mM glucose and 50 μM [U-^13^C]-succinic acid (Cambridge isotope CLM-1571-0.1). This buffer was pre-equilibrated at 37°C, 21% O_2_, and 5% CO_2_ prior to incubations. We incubated freshly dissected retinas in 200 μL media and eyecups in 100 μL, all at 37°C, 21% O_2_, and 5% CO_2_. To determine metabolite uptake or export rate we samples incubation media before (0 minutes), 20 minutes, or 40/45 minutes after the beginning of the incubation. Following incubations, media samples and tissue were flash frozen in liquid N_2_.

### Metabolite Extraction

Metabolites were extracted using 80% MeOH, 20% H_2_O supplemented with 10 μM methylsuccinate (Sigma, M81209) as an internal standard to adjust for metabolite loss during the extraction and derivatization procedures. The extraction buffer was equilibrated on dry ice, and 150 μL was added to each sample. Tissues were then disrupted by sonication and incubated on dry ice for 45 minutes to precipitate protein. Proteins were pelleted at 17,000 x g for 30 minutes at 4°C. The supernatant containing metabolites was lyophilized at room-temperature until dry and stored at -80°C until derivatization. The pellet containing protein was resuspended by sonication in RIPA buffer (150 mM NaCl, 1.0% Triton X-100, 0.5% sodium deoxycholate, 0.1% SDS, 50 mM Tris, pH 8.0) and the amount of protein was determined by a BCA assay (ThermoFisher, 23225).

### *Metabolite* Derivatization

Lyophilized samples were first derivatized in 10 μL of 20 mg/mL methoxyamine HCl (Sigma, Cat#: 226904) dissolved in pyridine (Sigma, Cat#: 270970) at 37°C for 90 minutes, and subsequently with 10 μL tert-butyldimethylsilyl-N-methyltrifluoroacetamide (Sigma, Cat#: 394882) at 70°C for 60 minutes.

### Gas Chromatography-Mass Spectrometry

Metabolites were analyzed on an Agilent 7890/5975C GC-MS using selected-ion monitoring methods described extensively in previous work (Du et al., 2015). Peaks were integrated in MSD ChemStation (Agilent), and correction for natural isotope abundance was performed using the software IsoCor (Millard et al., 2019). Corrected metabolite signals were converted to molar amounts by comparing metabolite peak abundances in samples with those in a ‘standard mix’ containing known quantities of metabolites we routinely measure. Multiple concentrations of this mix were extracted, derivatized, and run alongside samples in each experiment. These known metabolite concentrations were used to generate a standard curve that allowed for metabolite quantification. Metabolite abundance was normalized to tissue protein concentration, and following this, paired tissues such as retinas and eyecups from the same mouse were treated as technical replicates and averaged.

### Ex vivo oxygen consumption

Following euthanasia, mouse tissues were dissected and cut into quarters in Hank’s buffered salt solution. These tissues were incubated in Krebs-Ringer bicarbonate buffer (KRB) supplemented with 5 mM glucose and pre-equilibrated at 37°C and 5% CO_2_. We determined OCR using a custom-built perifusion flow-culture system (Neal et al., 2015; Sweet et al., 2002). Tissues were perifused in chambers between Cytopore beads (Amersham Biosciences, Piscatawy, NJ) and porous frits. With KRB supplemented with 5 mM glucose, 1x Antibiotic-Antimycotic (Gibco), and 1 mg/mL fatty acid-free bovine serum albumin. An artificial lung oxygenated supplemented KRB with a mixture of 21% O_2_, 5% CO_2_, and 74% N_2_. Oxygenated media was passed through a bubble trap before exposure to mouse tissues. Outflow media came into contact with a glass wall coated with a thin layer of oxygen sensitive polymerized Pt(II) Meso-tetra(pentafluorophenyl)porphine dye (Frontier Scientific, Logan, UT) painted on the inner glass wall of the chamber. Following a 405 nm light pulse, the dye-coated glass emits a phosphorescent signal detected at 650 nm. The decay lifetime is dependent on oxygen tension. The flow rate of KRB along with the quantitative relationship between dye phosphorescent decay and oxygen concentration were used to determine tissue OCR. All OCR measurements were obtained under control conditions (baseline, 5 mM glucose), one or more experimental conditions, and a ‘zeroed’ condition wherein 3 mM potassium cyanide (KCN) was used to directly inhibit complex IV and thus subtract the effect of residual non-mitochondrial oxygen consumption from our measurements.

### Lactate dehydrogenase assays

We quantified LDH levels using a CyQuant™ Cytotoxicity Assay kit (ThermoFisher C20300) and followed the manufacturers protocol. To obtain samples we dissected retinas and eyecups, incubated them in 200 μL (retina) or 100 μL (eyecup) KRB buffer, supplemented with 5 mM ^12^C-glucose and 50 μM ^13^C-succinate and indicated concentrations of digitonin. We sampled 5 μL of incubation media at the indicated times for analysis. At the end of the experiment we homogenized tissue in 200 μL of lysis buffer (10% Triton-X100) and 5 μL was used for tissue LDH quantification. We confirmed that all absorbance (490 nm) measurements were in the linear range of the reaction using a BioTek Synergy 4 plate reader.

### Radiation

To measure of permeability we compared volume of distribution of an extracellular marker (^14^C-sucrose) to of ^3^H_2_O inside and outside tissue. Uptake of 3H-H2O and 14C-sucrose (Perkin Elmer, Waltham, MA) was measured as carried out previously ^17^. ¼ retina or 1 eyecup was incubated in 200 μL KRB with 5 mM bicarbonate in 12 × 75 mm polypropylene test tubes at 37 degrees C for 15 min in a shaking water bath. We added 0.5 mCi ^3^H and 0.2 mCi for ^14^C to the tubes and incubated tissue in radiotracer for 60 min. Accumulation of radiolabel was determined by separating the cell-associated radioactivity (CAR) from the free radioactivity by transferring the cell suspension to a 0.4 mL centrifuge tube (USA Scientific, Ocala, FL) with a layer of 1:37.5 n-dodecane:bromo-dodecane (Sigma–Aldrich, St. Louis, MO) and spinning at 12,535 x g, 8 s. The tube was placed briefly in liquid N_2_ and we used a razer blade to cut through the radioactive-free oil layer. The bottom portion of the tube containing the tissue was placed into a 7 mL glass scintillation vial. We added 5mL of Ecolume liquid scintillation cocktail (MP Biomedicals, Cat No. 0188247001) per tube, vortexed the vials, and counted ^3^H and ^14^C using a Beckman liquid scintillation counter (Model LS6500). The CAR data for each sample was normalized by subtracting the non-specific values and then dividing by the dose. Quench correction for samples was not needed since counts were normalized to the dose and quenching did not vary from sample to sample.

### Statistical Analysis

We performed all statistical data analyses using Prism Version 8 (GraphPad Software). The significance threshold was set at p<0.05. To fit curves of oxygen consumption as a function of [succinate], for each sample we averaged steady-state oxygen consumption over >5 minutes at the end of a given treatment. These averaged values were considered to be the OCR at each given [succinate] for each sample. We fit the curve to an allosteric sigmoidal shape *(Figure1)*

## Funding Support

This research was supported by funding from the following sources T32 EY007031/EY/NEI NIH HHS/United States (to DTH), F31 EY031165/EY/NEI NIH HHS/United States (to CMB), R01 EY006641/EY/NEI NIH HHS/United States (to JBH), R01 EY017863/EY/NEI NIH HHS/United States (to JBH). OCR studies were performed at the Cell Function Analysis Core of the University of Washington Diabetes Research Center, supported by P30 DK017047/DK/NIDDK NIH HHS/United States.

## Competing Interest Statement

The authors declare that they have no conflicts of interest with the contents of this article.

## Notes

### Competing Interest Statement

The authors have declared no competing interest.

### Summary of Updates

Edited authors Edited language Edited figures

